# Competitive membrane wetting of polymer blends in artificial cells initiates phase separation and promotes fractionation

**DOI:** 10.1101/2022.03.23.485531

**Authors:** Chiho Watanabe, Tomohiro Furuki, Yuki Kanakubo, Fumiya Kanie, Keisuke Koyanagi, Jun Takeshita, Miho Yanagisawa

## Abstract

Biomolecular condensates driven by liquid–liquid phase separation (LLPS) have received attention as novel activity regulators of living organisms. In intracellular LLPS, an important question is what type of biomolecules form condensates under what conditions. In this regard, possible interactions between biomolecules have been investigated. Recently, LLPS condensates have been reported to regulate the membrane structure upon wetting. However, the possibility of membrane wetting, in which the membrane conversely regulates the LLPS, remains unexplored. Using droplets of short polyethylene glycol and long dextran blends encapsulated with a lipid membrane, we demonstrate that membrane wetting regulates LLPS in cell-size spaces and alters the equilibrium state. In smaller droplets, the two-phase region expands beyond the bulk system, and the fractionation degree increases, particularly during the separation between short PEG and long dextran. We explain the space-size dependent LLPS based on the competitive membrane wetting between the polymers. Smaller droplets promote the membrane wetting of short PEG, which enhances the depletion force between long dextran molecules and finally induces LLPS. This shows that competition for membrane wettability among various molecules can regulate LLPS in cell-size spaces, rendering this LLPS principle feasible in living cells.

## Introduction

In living organisms, liquid-liquid phase separation (LLPS) has emerged as a fundamental mechanism for regulating biological activity through biomolecular condensate formation. In intracellular LLPS, the type of biomolecules that separates to form condensates and the conditions under which this occurs are to be determined. LLPS-forming biomolecules can be identified via the molecular-level analysis of condensates using nuclear magnetic resonance spectroscopy, small-angle X-ray scattering,etc.^1, 2, 3^ Meanwhile, theoretical researchers are attempting to clarify the conditions for initiating the LLPS of biomolecules by calculating the possible interactions between various molecules, including solvents and ions.^4, 5, 6^ Previous studies regarding LLPS focus on molecules in the cytoplasm and rarely consider the contribution of membrane structures.

Recently, Agudo-Canalejo et al. have reported the role of membranous structures known as autophagosomes for the removal of LLPS condensates in the autophagy process.^7^ The membrane wetting of LLPS condensates facilitates in the emergence and disappearance of autophagosomes. Such a wetting effect of LLPS condensates on the membrane has been previously reported in LLPS studies where liposomes are used as artificial cells prior to cells. For example, when a polyethylene glycol (PEG)/dextran blend undergoes LLPS in liposomes, the PEG/dextran interface and PEG membrane adsorption deform the liposome.^8, 9^ These studies suggest a new role for LLPS condensates in cells via membrane wetting, namely, the regulation of membrane structures.

Contrary to the wetting effect of LLPS condensates on membranes, the wetting effect of membranes on LLPS is rarely reported. Previously, we evaluated the membrane effect on molecular behaviors in cell-size spaces using water-in-oil (W/O) droplets encapsulated with a lipid monolayer; it is revealed that the effect can be enhanced by changing the membrane lipid composition and increasing the surface-to-volume ratio of the droplets.^10^ Furthermore, we discovered acceleration in LLPS domain coarsening in small droplets, as compared with the bulk for PEG/DNA^11^ and PEG/gelatin^12^ blends in a two-phase region distant from the critical point. Meanwhile, the equilibrium state did not change significantly. The enhanced membrane effect changes various phenomena in 1-phase solutions, such as nanostructure transition^13, 14^, gene expression,^10^ and molecular diffusion^15, 16, 17^. Therefore, the enhanced membrane wetting effect on LLPS in cell-size spaces is likely to appear in the 1-phase region near the critical point that determines the condition of LLPS emergence. The investigation of such a near-critical point is important for understanding intracellular LLPS, where the emergence/disappearance of condensates is repeated in response to intracellular activity^18^

In this study, we used W/O droplets containing a 1-phase binary polymer blend in bulk to analyze the membrane wetting effect on LLPS. It is advantageous to use droplets as molecular exchange through the membrane is negligible, and membrane wetting is not easily converted into membrane deformation until the solution reaches phase equilibrium. Liposomes without these features deform their membranes via dehydration, depending on the inner solution.^19, 20^ As a polymer blend, we selected the PEG/dextran blend because it is one of the most popular polymer blends with LLPS, and the wetting effect of LLPS on the liposome membrane has been reported.^19, 20^ For cell-sized droplets with a radius *R* < 20 μm, we discovered that the critical point of the PEG/dextran blend shifts to a lower concentration, and that the two-phase region expands over the bulk system. In addition, a decrease in *R* alters the equilibrium composition and increases the fractionation of short PEG and long dextran. We explain this *R*-dependent LLPS by the preferential membrane wetting of PEG and depletion force between the large dextran molecules, which is enhanced in a small space. The results demonstrate that competition for membrane wettability between polydisperse molecules is essential to the separation of molecules in cell-size space. This polydisperse environment can be a prototype of the intracellular environment, suggesting that the membrane structures and cytoplasmic components cooperate to control the appearance and disappearance of LLPS condensates. This may provide important insights for understanding the dysregulation mechanism of LLPS, which has been suggested to trigger diseases.^21^

## Results

### Cell-sized space initiates phase separation of PEG/dextran blend

The PEG/dextran blend is known to separate into two liquid phases (2-phase) depending on the polymer composition and temperature. First, we determined the phase diagrams of binary polymer blends of PEG with a molar mass of 6 kg/mol (PEG6k) and dextran with a molar mass of 450-650 kg/mol (dextran500k) in bulk at approximately 20 °C via turbidity confirmation and microscopic observation (Figure 1a). The boundary separating the 2-phase region (pink) from the 1-phase region (white) coincides with the binodal line obtained from four different compositions with PEG/dextran = 1:1 (solid orange line with open orange diamonds in Figure 1a; see also Figure S1). The dotted gray lines represent tie lines. Mixing the 2-phase solution via vortexing rendered the solution opaque, as shown by the solution of 5 wt% PEG and 5 wt% dextran (Figure 1b). The overall phase diagram and critical concentration (approximately 4 wt% PEG6k and 4 wt% dextran500k) were similar to those of previous relevant studies involving the use of PEG with a molar mass of 8 kg/mol and dextran500k.^22, 23, 24^

**Figure 1.**
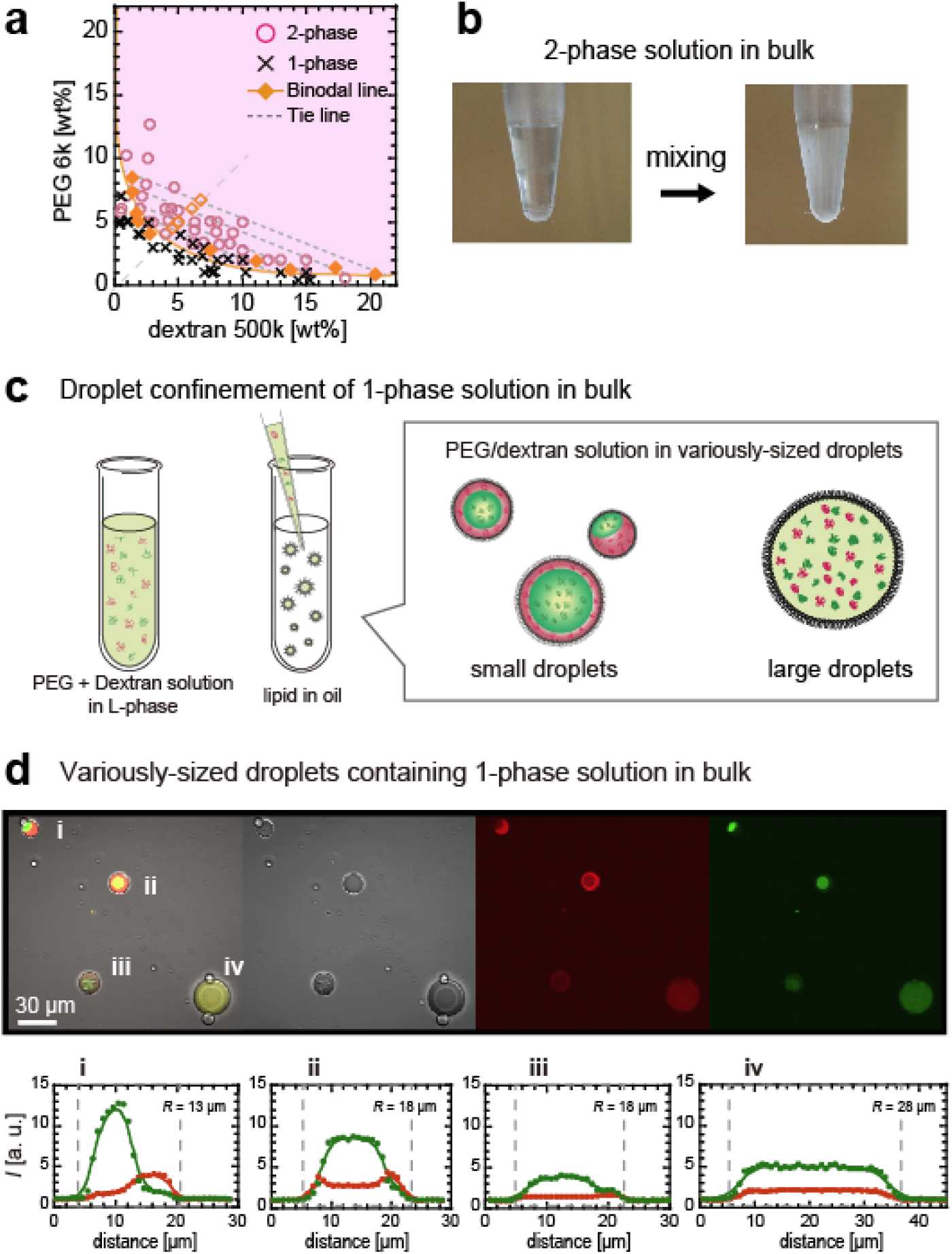
(a) Phase diagram of PEG6k and dextran500k blends in bulk. Two- and 1-phase regions are shown in pink (circle) and white (cross), respectively. Binodal line and tie lines derived from four different compositions (PEG6k/dextran500k = 1:1, diamond) are indicated by solid orange line and dotted gray line. (b) Bulk solution of 5wt% PEG6k and 5wt% dextran500k in two-phase becomes opaque when mixed with vortex. (c) Illustration of PC droplet preparation containing PEG/dextran blend. (d) (top) Confocal images of PC droplets containing 1-phase solution in bulk, 3 wt% PEG6k, and 3wt% Dextran500k. (From left to right) Merged image of transmission image and fluorescent images of RB-PEG5k (red) and FITC-Dex500k (green). (bottom) Fluorescence intensity *I* profile of RB-PEG5k (red) and FITC-Dex500k (green) along equatorial plane of droplets. Dotted lines indicate droplet surface positions. Radius of droplets is 13 μm (i), 18 μm (ii, iii), and 28 μm (iv).

In this study, all experiments were performed using 1-phase solutions in bulk (white region in Figure 1a). To investigate the membrane wetting effect, we prepared PEG/dextran droplets encapsulated with a phosphatidylcholine (PC) lipid layer (Figure 1c). The droplet radius *R* ranged from 5 to 70 μm, and the smaller droplets had a larger surface-to-volume ratio, resulting in a more effective membrane wetting system. Surprisingly, the PEG/dextran blend in small droplets transited from 1-phase to 2-phase, whereas that in large droplets with *R* >> 20 μm remained in 1-phase. Figure 1d (top) presents an example of a microscopic image showing such *R*-dependent phase separation, where a solution of 3 wt% PEG and 3 wt% dextran blends was confined in droplets with different radii (*R* = 13–28 μm. For fluorescence observation, 0.1 mol% of rhodamine-B labeled as PEG5k (RB-PEG5k) and fluorescein isothiocyanate labeled as dextran500k (FITC-Dex500k) were added to the original PEG6k and dextran500k solutions, respectively. For small droplets with *R* < 20 μm (i-iii), clear (i, ii) or slight (iii) phase separations were observed, where the PEG-rich and dextran-rich phases are shown in red and green, respectively. The fluorescence intensity profiles suggested that the fractionation degree increased with *R* (Figure 1d, bottom). By contrast, the larger droplet with *R* = 28 μm (iv) indicated no phase separation, as in the bulk.

We formed two hypotheses based on these findings: First, for small droplets, the 1-phase solution in bulk phase separates; hence, the 2-phase region expands beyond the bulk system. Second, the fractionation degree of the phase-separated polymers increases with the droplet size. To verify these hypotheses, we used droplets of various sizes in our investigation.

### Phase equilibrium states based on droplet size

The 1-phase solutions of PEG6k/dextran500k in bulk (marked in white in Figure 1a) had likely transitioned to a 2-phase region inside the small droplets. To clarify such a 2-phase region, we classified the 2-phase region into two categories based on the droplet radius *R*: (i) The 2-phase region in all droplets of any *R*, and (ii) the 2-phase region in small droplets (Figure 2a, left). The 2-phase region for any *R* droplet (i) corresponds to the 2-phase region in the bulk (pink, Figure 1a). Hereinafter, we refer to (i) as the 2-phase region.

**Figure 2.**
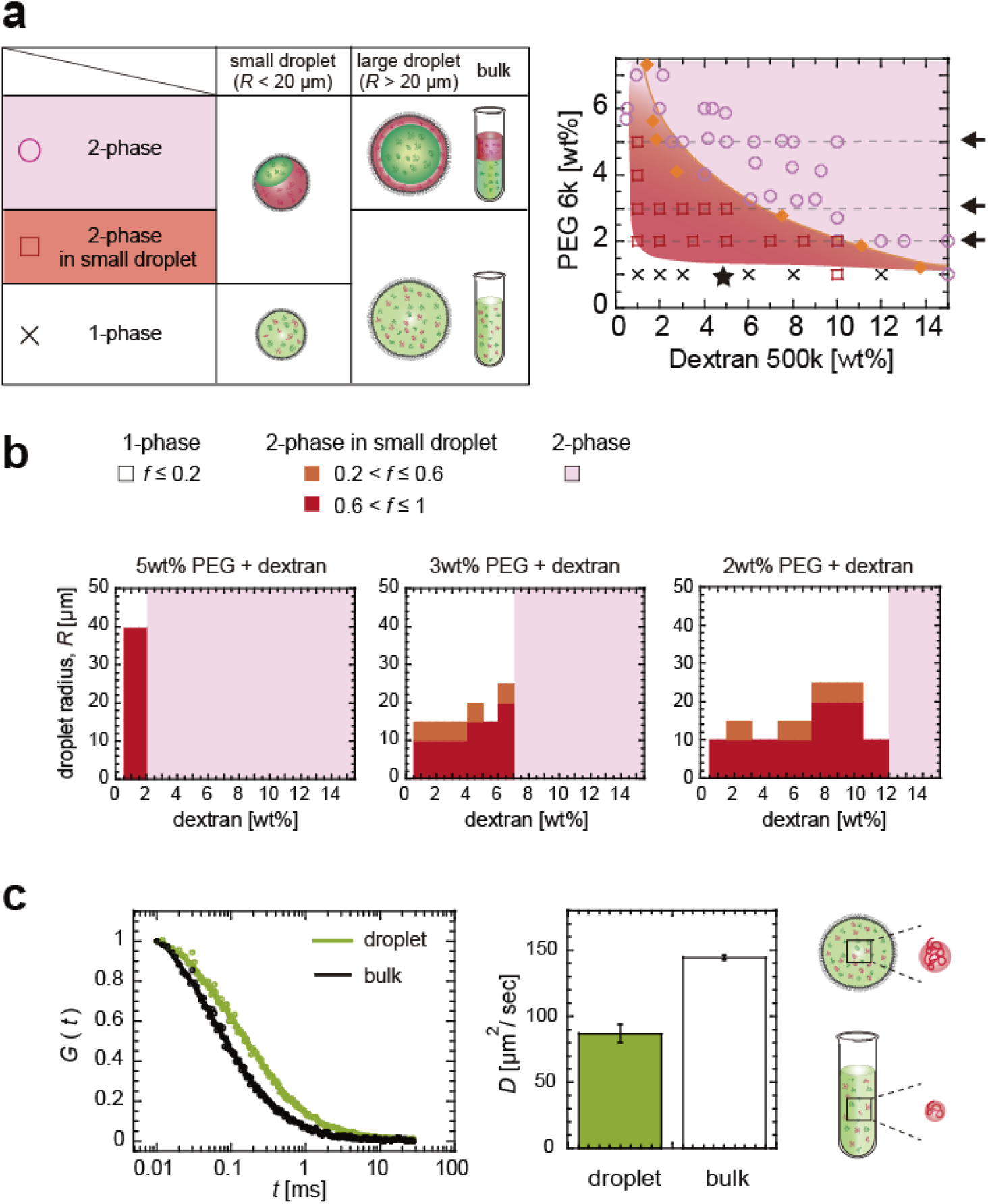
(a) Phase diagram of PEG6k and dextran500k blends in droplets with radius *R* ≥ 5μm. In 2-phase region (pink, circle), droplets of any *R* phase separate, which is likely due to bulk system, unlike in 1-phase region (white, cross). *R*-dependent phase separation occurs in 2-phase region in small-droplet region (red, square) (b) Relationship between three phase regions and *R*. Dextran concentration is varied under fixed concentrations of PEG (from left to right, 5 wt%, 3 wt%, and 2 wt%, indicated by arrows in Fig. 2a). Two-phase region in small droplets (red and orange) and 1-phase (white) regions are defined based on frequency of phase-separated droplets, *f*. (c) Molecular diffusion of 40 nM RB-PEG5k in 1-phase region, as well as those of 1 wt% PEG and 5wt% dextran500k (marked with a star in Fig. 2a). (left) Autocorrelation function G(τ) in bulk (black) and in droplet with a radius *R* ~7 μm (green). (right) Diffusion coefficient *D* for small droplets with a radius *R* = 17.5 ± 9.0 μm (Ave. ± SD.; n = 18) and for bulk (n = 19).

Figure 2a (right) shows the phase diagram for droplets with a size of 5 μm ≤ *R* ≤ 50 μm, where the two different 2-phase regions, (i) and (ii), and the 1-phase region are marked in pink (circle), red (square), and white (cross), respectively. This clearly demonstrates that the 1-phase region in the droplet decreases compared with that in the bulk (white, Figure 1a) owing to the appearance of the 2-phase region in the small droplets (marked in red). In addition, the critical point appeared in a lower concentration solution, *i.e.*, instead of in PEG 4wt% and dextran 4wt%, it appeared in PEG 2wt% and dextran 1wt%.

To determine the upper limit *R*\* of the droplet size and the composition at which the PEG/dextran blends transition from the 1-phase to the 2-phase region inside the droplets, the frequency *f* of the phase-separated droplet at a droplet size width of 5 μm was analyzed by changing the dextran concentration *C*_dex_ under a fixed PEG concentration (2 wt%, 3 wt%, and 5 wt%, indicated by arrows in Figure 2a). In terms of *f*, we identified the 1-phase region as *f* ≤ 0.2 (white), 2-phase region in the small droplet as *f* > 0.6 (red), and the intermediate region as 0.2 < *f* ≤ 0.6 (orange) (Figure 2b). At all the PEG concentrations evaluated, the 2-phase region in the small droplet (ii) was observed at *C*_peg_ ≥ 1 wt%, which was lower than the bulk values (2-phase (i), marked in pink). Although the *R*\* for *C*_peg_ = 5 wt% was approximately 40 μm, the *R*\* for *C*_peg_ = 2 and 3 wt% was approximately 20 μm, regardless of *C*_dex_.

In the 1-phase region (white, Figure 2a), fluorescence microscopy observations showed no distinguishable difference between the droplet and bulk systems. Because a difference might be indicated below the optical resolution, we analyzed the diffusion coefficient *D* of RB-PEG5k for 1 wt% PEG6k and 5wt% dextran500k, which was in the 1-phase region in the bulk (marked with a star in Fig. 2a). We derived *D* for small droplets with *R* = 17.5 ± 2.1 μm (Ave. ± SD., n = 18) and compared with that in the bulk (Figure 2c) by analyzing the autocorrelation functions using fluorescence correlation spectroscopy (FCS, see the Method section). The *D* for the small droplet and the bulk was 86.8 ± 6.7 μm^2^/s (n = 18) and 144.3 ± 1.8 μm^2^/s (n = 19), respectively. The smaller *D* of the droplets than that of the bulk suggests that the 1-phase solution inside the droplets contains molecular clusters comprising two or three molecules of PEGs, although the cluster size might not be sufficiently large to initiate nucleation growth. Under macromolecular crowding conditions, the *D* of the small droplets is lower than that of the bulk ^15, 16, 17^ and transient cluster formation^25, 26, 27^ have been reported from simulation and experimental studies for various protein and polymer systems in the 1-phase region. Therefore, microscopic nonuniformity can be a typical condition for semi-diluted or condensed polymer solutions.

These data strongly support one of our hypotheses: the cell-sized space shifts the critical point to a lower concentration and expands the 2-phase region beyond the bulk system. Even in the remaining 1-phase region, a few PEG molecules may form clusters.

To establish the phase equilibrium state in the small droplets, the fractionation degree was analyzed for the phase-separated droplets. As shown in Figure 3a (left), RB-PEG5k (red) and FITC-Dex500k (green) separated into two phases, and no visible localization to the droplet surface was observed. Assuming that the fluorescence intensities of RB-PEG5k and FITC-Dex500k were proportional to the respective concentrations of unlabeled PEG6k/dextran500k, as reported previously^22^, we evaluated the fractionation degree via intensity analysis. Based on the average intensity of RB-PEG5k and FITC-Dex500k for the dextran-rich and PEG-rich phases, we obtained the *I*_max_ and *I*_min_ of RB-PEG5k (or FITC-Dex500k) for the PEG-rich and dextran-rich phases (or dextran-rich and PEG-rich phases), respectively (Figure 3a, right). The intensity difference normalized by the background, *ΔI/I*_0_, where *ΔI* = *I*_max_ – *I*_min_, was analyzed for various compositions of the 2-phase region in the small droplet. The PEG concentration was fixed at 3 wt% and 2 wt%, and the dextran concentration was varied (indicated by the arrows in Figure 2a). The derived values of *ΔI*/*I*_0_ were plotted against *R* (Figure 3b). For all the polymer compositions evaluated, the values of *ΔI/I*_0_ for RB-PEG5k and FITC-Dex500k increased as *R* decreased to less than ~20 μm. These data support the hypothesis presented in Figure 1, i.e., droplets with a *R*\* smaller of ~20 μm have a greater fractionation degree. This implies that the Gibbs energy, which determines the phase equilibrium state, decreases in a droplet size-dependent manner below *R*\*.

**Figure 3.**
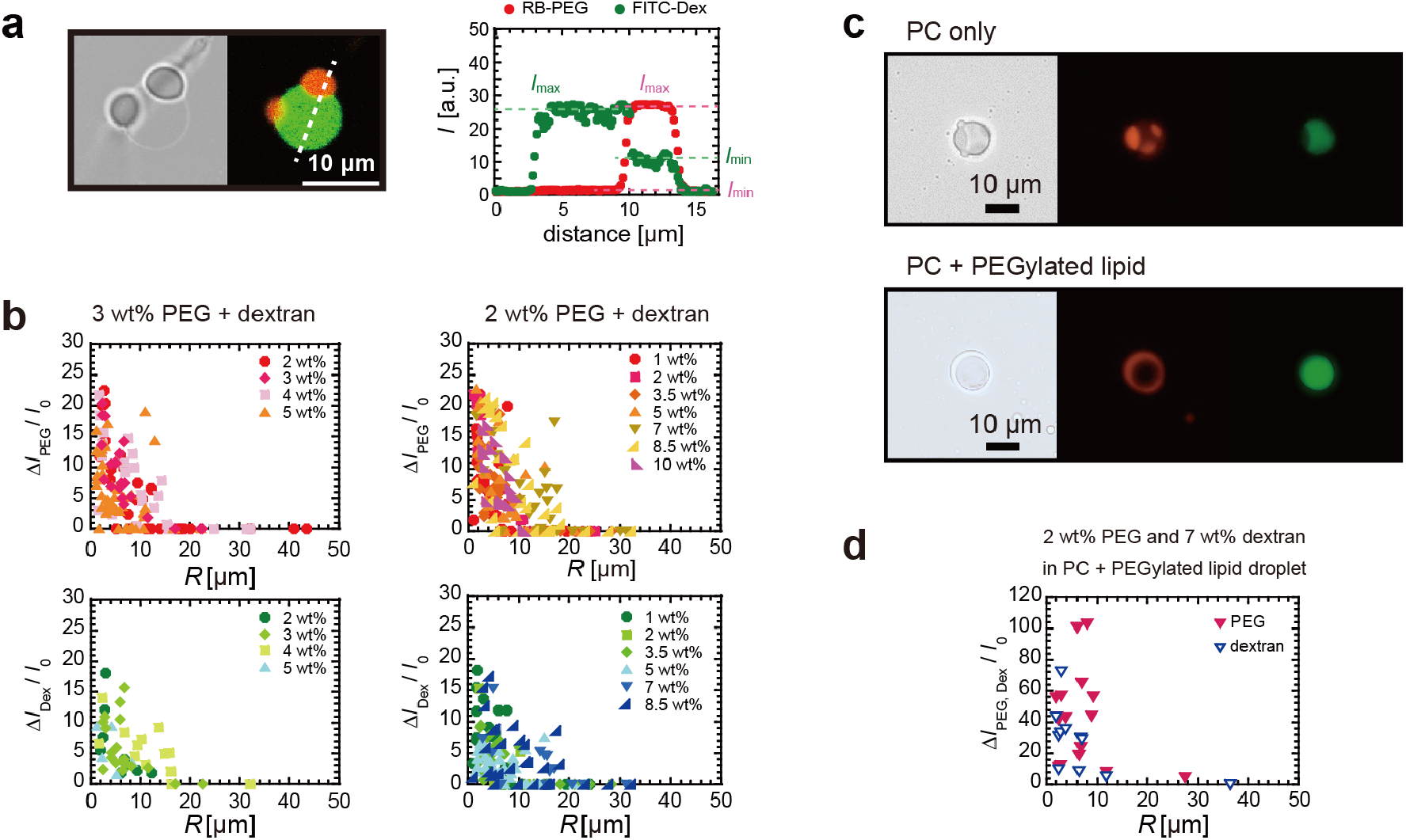
(a) Microscopic image of PC droplet containing 2 wt% PEG and 7wt% dextran with RB-PEG5k (red) and FITC-Dex500k (green). Fluorescence intensity profile shown along broken line. Fractionation degree evaluated as normalized intensity difference, *ΔI/I_0_* where *ΔI* = *I*_max_ *I*_min_ in the droplet of interest and *I*_0_ is the background intensity. (b) *R* dependence of *ΔI*/*I*_0_ of RB-PEG5k (upper) and FITC-Dex500k (bottom) for fixed PEG concentration to be 3 wt% (left) and 2 wt% (right) against various dextran concentrations. (c) Transmission and fluorescence images of 2 wt% PEG (red) and 7 wt% dextran (green) inside droplets encapsulated with PC only (upper) and PC with 25 mol% PEGylated lipids (lower). (d) *R* dependence of *ΔI/I_0_* for RB-PEG5k (red) and FITC-Dex500k (blue) with 2 wt% PEG and 7wt% dextran, respectively, inside droplets encapsulated with PC containing 25 mol% PEGylated lipids.

PEG has been reported to adsorb the PC bilayer membrane,^8, 28^; hence, the membrane wetting of PEG might have triggered the phase separation of the PEG/dextran blends inside the PC droplets. To enhance the membrane wetting effect of PEG, 25 mol% PEGylated lipids were added. In the early stages, complete wetting to the PEG-rich domains was observed more frequently in PEGylated lipid droplets compared to PC droplets (Figure 3c). In addition, the value of *ΔI/I*_0_ for RB-PEG5k was higher than that for the PC droplets (Figure 3d). Furthermore, the PEGylated droplets accelerated significantly, as compared with the PC droplets (Supplemental Figure S2). These data clearly show that membrane wetting is enhanced by the addition of PEGylated lipids, as expected, and that the preferential membrane wetting of PEG is key in initiating *R*-dependent phase separation. By contrast, the critical size *R\** is comparable for PC-only and PC with PEGylated lipid systems (~20 μm). This implies that a further increase in the membrane wettability for PEG changes the equilibrium polymer composition but does not significantly change the critical point, which determines the LLPS emergence condition.

### Distribution of polydisperse dextran inside phase-separated droplets

*R*-dependent phase separation inside the droplets was observed for 1-phase solutions in bulk with a low polymer concentration. Under such a low concentration of PEG8k/Dex500k blends, Liu et al. discovered that the Mw of dextran in the dextran-rich phase was significantly higher than that in the coexisting PEG-rich phase. By contrast, PEG in the two coexisting phases indicated similar Mw between the coexisting 2-phase regions.^8^ In our system using PEG6k/dextran500k blends, a difference in the Mw of dextran between the coexisting PEG-rich and dextran-rich phases is expected. If such a heterogeneous spatial distribution of polydisperse dextran500k is realized, then the longer dextran with a higher Mw is expected to be partitioned into the dextran-rich phase in the small droplets, instead of the shorter dextran. To test this hypothesis, we analyzed the spatial distribution of dextran with a Mw lower than the original Mw of 500k using FITC-Dex150k and FITC-Dex10k. The radius of gyration of dextran150k (~10 nm) is smaller than that of dextran500k (~17 nm) but larger than that of dextran10k and PEG6k (~3 nm).

For the 3 wt% PEG6k and 3 wt% dextran500k blends, the partition of FITC-Dex150k and the *R* dependence of *ΔI/I*_0_ are shown in Figure 4a; as shown, they were similar to that of FITC-Dex500k (Figures 1–3). Although FITC-Dex10k was partitioned in the dextran-rich phase, the *R* dependence of *ΔI/I*_0_ was lower compared with those of FITC-Dex150k and FITC-Dex500k, thereby resulting in a broader *ΔI/I*_0_ distribution against *R* (Figure 4b). The observed tendency pertaining to the equilibrium polymer composition in the droplet system, *i.e*., the stronger partitioning of higher-Mw dextran in the dextran-rich phase, is consistent to that near the critical point in the bulk system.^8^ This heterogeneous distribution of polydisperse dextran500k may be related to the membrane wetting of PEG, which has been suggested as the contributor to R-dependent phase separation within PC droplets.

**Figure 4.**
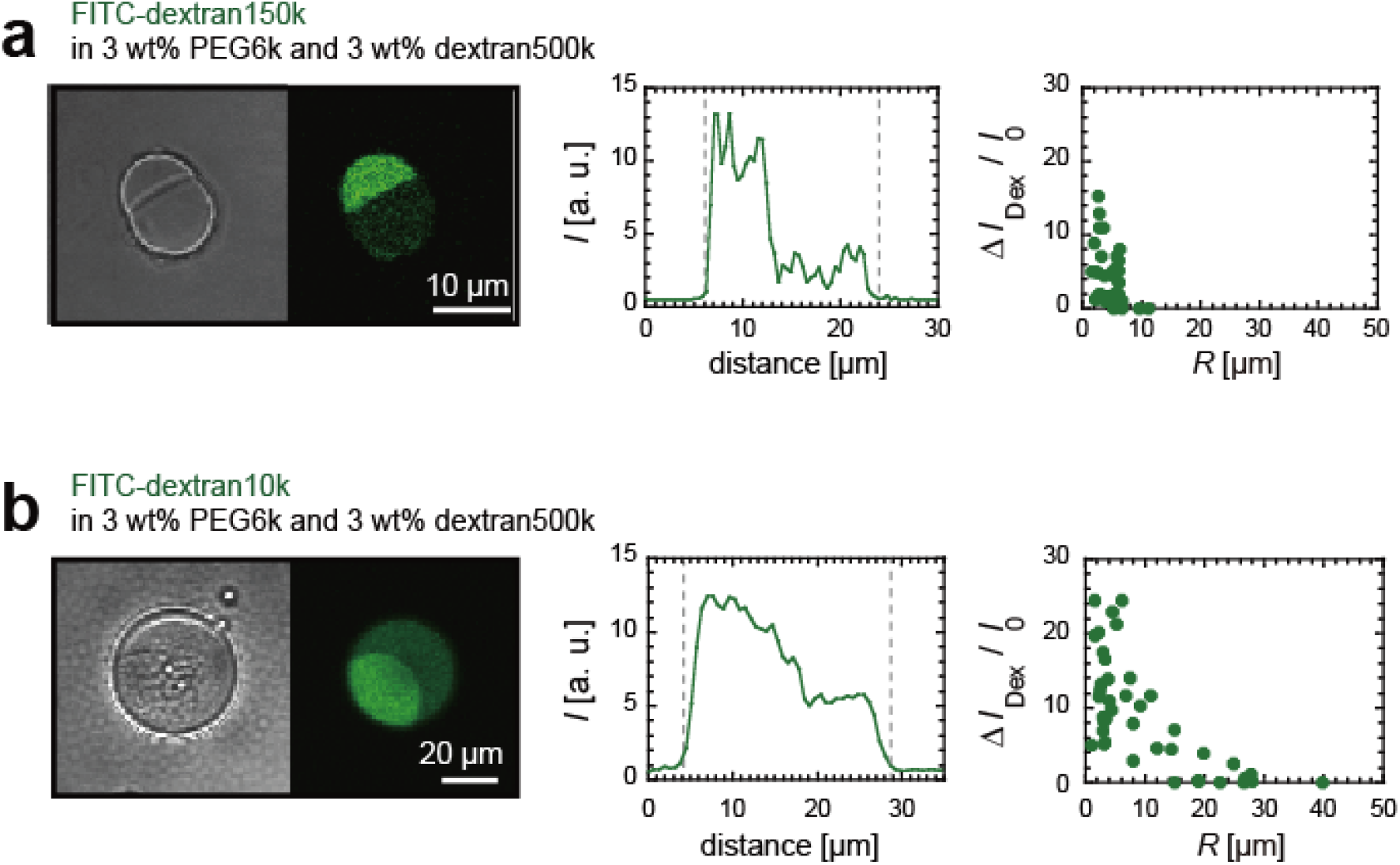
(left) Microscopic transmission and fluorescence images of PC droplet containing 3 wt% PEG and 3wt% dextran with 0.1 mol% FITC-Dex (green) of Mw = 150k (a) and 10k (b). (center) Fluorescence intensity profile along equational plane. (right) Normalized intensity difference, *ΔI/I*_0_, between PEG-rich and dextran-rich phases plotted against droplet radius, *R*.

### Competitive membrane wetting between PEG and polydisperse dextran

We discovered that the membrane wetting of PEG and the heterogeneous distribution of long and short dextran molecules were crucial for understanding the mechanism underlying *R*-dependent phase separation. To evaluate the membrane wettability between PEG and polydisperse dextran, we analyzed the surface tension γ of the PC droplets using the pendant drop method. The values of γ (average (Ave.) ± standard error (SD.)) for 2wt%, 3wt% dextran500k, and 2 wt% dextran10k were 30.3 ± 2.9 mN/m (n = 30), 30.5 ± 1.4 mN/m (n = 26), and 20.3 ± 2.1 mN/m (n = 11), respectively (Figure 5a). The γ values for their mixture, 2 wt% dextran500k and 0.1 wt% dextran10k, were similar to that of 2 wt% dextran10k, i.e., 21.2 ± 4.3 mN/m (n = 15); in fact, this value is similar to that for the minor component, i.e., dextran10k.

**Figure 5.**
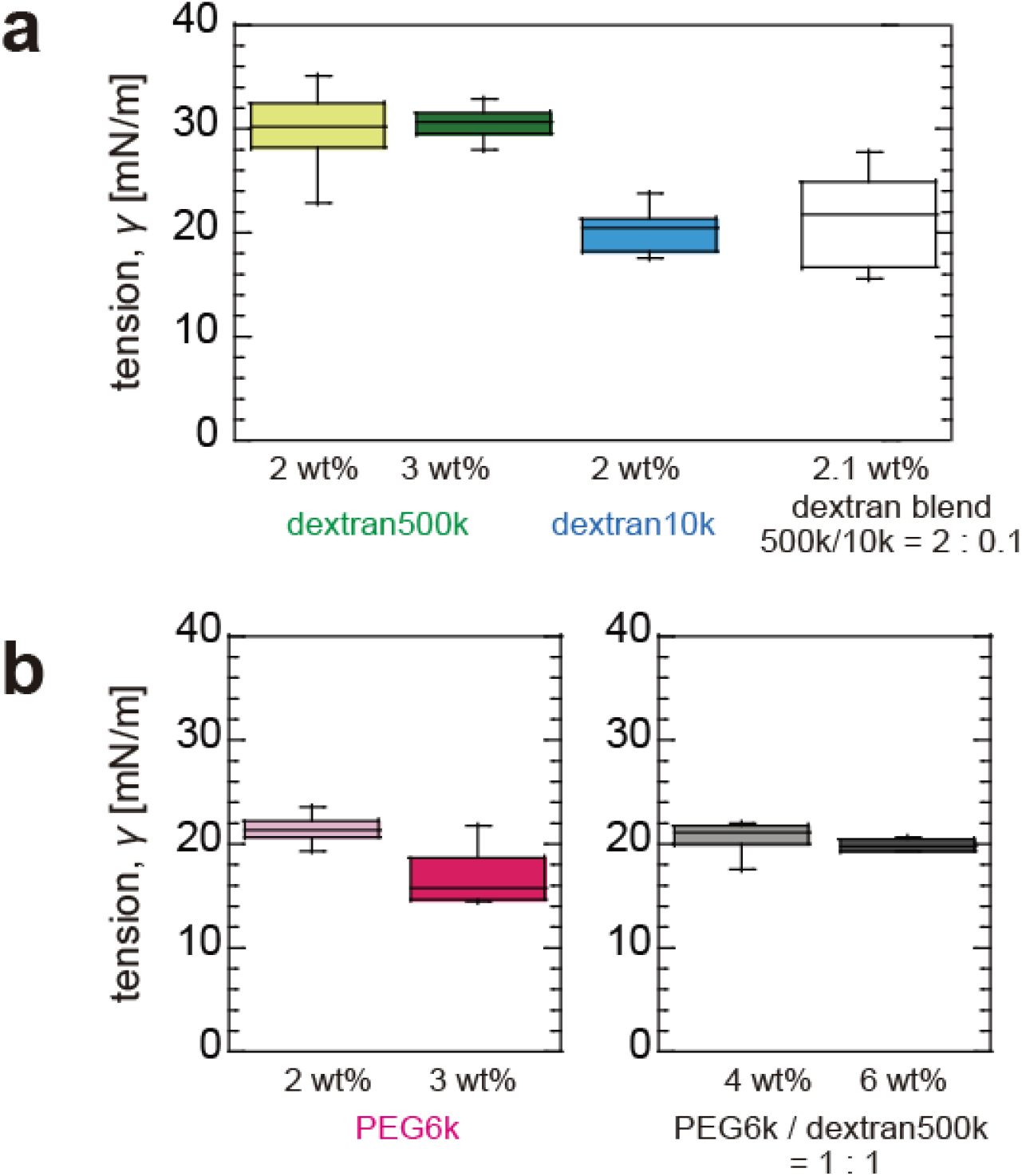
Surface tension of PC droplets containing different polymer solutions. (a) 2, 3 wt% dextran500k solutions, 2 wt% dextran10k solution, and blend of 2 wt% dextran500k and 0.1 wt% dextran 10k. (b) 2, 3wt% PEG6k solutions and blends of equal weight concentrations of PEG6k and dextran500k at 2wt% and 3wt%.

Meanwhile, the γ values for 2 wt% and 3 wt% PEG6k were 21.3 ± 1.1 mN/m (n = 25) and 16.9 ± 2.9 mN/m (n = 6), respectively (Figure 5b, left), where were similar to that for dextran10k and lower than for dextran500k. This trend is consistent with previous reports, thereby suggesting the higher membrane wettability of PEG compared with that of dextran with a higher Mw^8, 28^. The γ values for the PEG/dextran blends (total polymer concentration of 4 wt% and 6 wt%; PEG/dextran = 1:1 wt%) were 20.5 ± 1.8 mN/m (n = 18) and 19.9 ± 0.6 mN/m (n = 5), respectively (Figure 5b, right). These values are much lower than those of dextran500k (~30 mN/m) and similar to those of PEG and dextran10k systems (17–21 mN/m). Based on these data, we discovered that the surface tension for all the conditions evaluated did not depend significantly on the polymer concentration. This implies that the small PEG and dextran with Mw << 500k adsorbed preferentially to the membrane, thereby reducing membrane contact with the long dextran of lower wettability.

It is noteworthy that these γ values (17–30 mN/m) were obtained using millimeter-sized droplets. Therefore, they should be much higher than those of the micrometer-sized PC droplets. This is because such millimeter-sized droplets require a much longer time to reach the equilibrium value upon slow lipid adsorption to the W/O surface as compared with the micron-sized droplets.^29^ However, we could not analyze the γ for the micron-sized droplets containing PEG/dextran blends because the blends caused phase separation, and their viscoelasticity increased with time. To evaluate the possible artifacts arising from the slow relaxation at the droplet surface upon lipid adsorption, we performed similar experiments using the nonionic Span 80, which showed a much faster relaxation and a smaller γ of ~3 mN/m,^29^ as shown in Figure S3. An extremely similar *R* dependence in the partition of FITC-Dex10k in the dextran-rich phase and *ΔI/I*_0_ was observed in the Span 80 droplets. Therefore, we conclude that the relaxation process of lipids to micrometer-sized droplets is negligible in the *R*-dependent phase separation of the micron-sized droplets.

## Discussion

### Possible factors that induce macroscopic phase separation following membrane wetting

To address the mechanism by which the cell size space initiates *R*-dependent phase separation between short PEG6k and long dextran500k, we focus on the change in droplet surface energy due to preferential membrane wetting of PEG6k and shorter dextran with Mw << 500k. From Figure 5, the estimated change Δ*γ* by the adsorption of them for the micron-sized droplets is less than 10 mN/m. In addition, the surface energy gain increased in the smaller droplets with a larger surface-to-volume ratio. Therefore, we estimated the interfacial energy gain per unit volume at the droplet surface as follows:

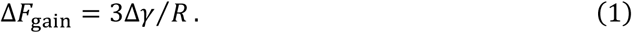

*ΔF*_gain_ was plotted against *R* for Δγ = 0.1, 1, and 5 mN/m (Figure 6a). When Δγ was 0.1 or 1 mN/m, *ΔF*_gain_ decreased abruptly below *R* ~20 μm. This implies that the adsorption of small PEG and shorter dextran molecules decreased the surface energy, particularly for the smaller droplets with *R* < 20 μm. Because this *R*-dependent and the size scale corresponds to an upper critical size *R*\* of ~20 μm (Figure 3b), the preferential wetting of small PEG and dextran over dextran500k is likely to be the origin of *R*-dependent phase separation in the cell-sized space.

**Figure 6.**
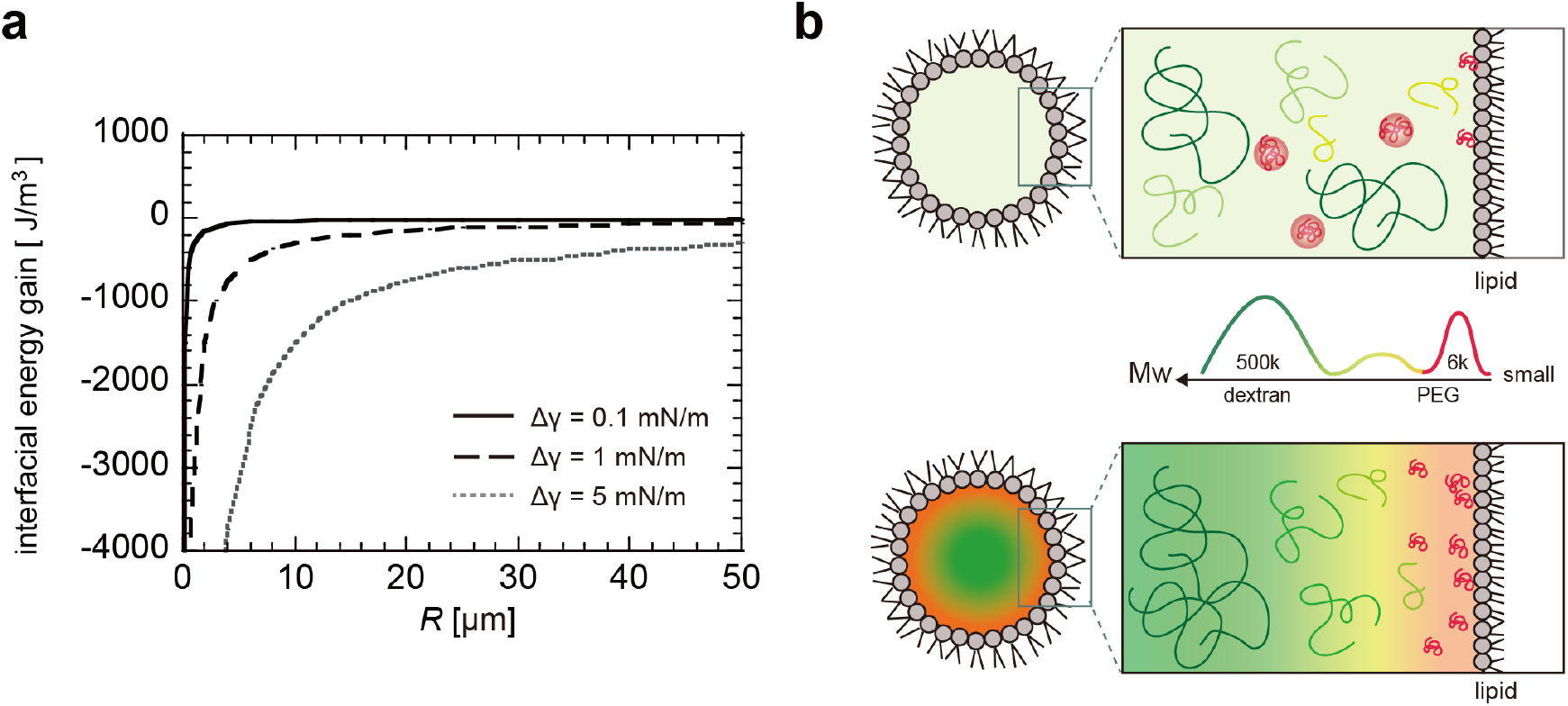
(a) Gain of interfacial energy per volume when surface tension is reduced by Δγ = 0.1 (solid line), 1 (dashed line), and 5 mN/m (dotted line), respectively. (b) Schematic illustrations of polymer blend of short PEG and polydisperse dextran inside artificial cells. (upper) Microscopic heterogeneity in 1-phase region; (bottom) Membrane-wetting-induced 2-phase region.

However, the membrane wetting effect is normally limited to the nanometer scale near the membrane surface. Therefore, other effects must be considered to explain the macroscopic phase separation inside the droplets following membrane wetting. If two or three molecules of PEGs exist as metastable nanometer-sized clusters beforehand inside the small droplets, preferential wetting of PEG and short dextran expects to initiate the macroscopic phase separation following membrane wetting (Figure 6b). In fact, slow diffusion of PEG inside cell-sized droplets suggests the presence of such metastable molecular clusters inside the small droplets (Figure 2c). In addition, this multi-step nucleation-type phase separation corresponds the nucleation process of prion-like LLPS in the bulk.^30^

Upon the membrane wetting, PEG6k and short dextran with a lower Mw appeared near the membrane covering the droplet surface, whereas the long dextran with a higher Mw localized at the droplet center, as illustrated in Figure 6b.^8, 31^ This implies that the compositional bias due to the membrane wetting can facilitate the phase separation of PEG/dextran blends near the critical point. This is because the heterogeneous distribution of polydisperse dextran (In Figure 4, short and long dextran molecules tend to localize in the PEG-rich and dextran-rich phases, respectively) corresponds to the equilibrium polymer composition near the critical point, as reported for PEG8k/dextran500k^8^.

In addition, the accumulation of small PEG on the membrane might enhance the effective depletion force between larger dextran molecules^32^. We estimated the depletion force exerted on the dextran500k molecules due to PEG6k (see Supplemental Information S1). For example, 3 wt% PEG6k (R_g_ ~3 nm) and 3 wt% dextran500k (R_g_ ~17 nm) corresponded to 5 mM and 60 nM, respectively. Assuming that the apparent radius of PEG6k was maintained as *R*_g_ and the sphere of *R*_g_ occupied the volume, the volume fraction of PEG6k was ~38 %, and the depletion force was estimated to be 3.5 *k*BT, which is larger than the binding force of PEG on the bilayer (~1.6 *k*_B_T^8^). Although the increase in PEG concentration slightly decreases the apparent size of the PEG chain^33^ and the effective volume fraction of PEG6k, the depletion force of ~3.5 *k*BT may be maintained or increased when the local concentration of PEG increases via membrane wetting. The short molecules contained in polydisperse dextran500k, (for example, *R*_g_ of dextran10k is similar to that of PEG6k) may also increase the depletion force between the large dextran molecules; however, the increase degree should depend on the fraction among the polydisperse dextran500k molecules and the amount of dextran accumulated on the membrane.

From a kinetic perspective, the viscoelastic property of the dextran-rich phase containing higher-Mw dextran may propel the small PEG and dextran molecules to the droplet surface owing to its elastic contraction, which is known as viscoelastic phase separation.^34^ This may correspond to the stepwise and slow increase in the PEG concentration upon phase separation, unlike the case of dextran (Figure S3, a). The faster adsorption of PEG compared with the condensation of dextran on the membrane may have contributed to the accelerated phase separation in the PEGylated lipid droplets (Figure 3S, b); this is because the rough dextran mesh does not interfere with PEG diffusion. This may correspond to the absence of isolated domains at the early stage of phase separation which is normally observed for the nucleation and growth.

### Mechanism of *R*-dependent phase separation in cell-size space

Herein, we propose a novel mechanism to explain the emergence of LLPS in small spaces via competitive surface wetting among the various molecules. The difference in the Mw-dependent surface wettability supports the preferential accumulation of small molecules on the surface. It triggers LLPS via three effects: 1) effectively decreasing the surface energy of the small system, 2) rendering the system composition more similar to the Mw-dependent equilibrium composition, and 3) enhancing the depletion force between large molecules. These effects should be applicable to various polydisperse small systems, where many small molecules with higher surface wettability are present along with a few large molecules with lower surface wettability.

Our polymer blend droplets encapsulated with a lipid membrane exhibit the structural characteristics of cells, i.e., polydisperse macromolecules in the micrometer-sized space of membrane structures. Therefore, it would be meaningful to verify whether the mechanism revealed herein is applicable to living cells, i.e., whether the membrane structures and cytoplasmic materials correspond to the emergence and disappearance of LLPS condensates, as well as to the composition of LLPS condensates via competitive membrane wettability among small and large molecules. Small molecules with high wettability that are not involved in the LLPS condensates behave as an inducer of LLPS. Therefore, it is necessary to analyze the size difference between the molecules in the LLPS condensates and the surrounding molecules, particularly near the membrane structures, as well as the affinity between these molecules and the membrane structures. Since the effective size and wettability of biomolecules in cells can change according to cellular activities, a multifaceted approach based on experimental research using cells and artificial cells as well as based on theory should be developed.

In summary, our findings are applicable to the thermodynamics of small polymer droplets and their application in food and biomedical materials. In addition, the LLPS mechanism revealed herein contributes to biophysics via the possible role of the membrane structures involved in LLPS via membrane wetting.

## Materials and Methods

### Materials

To prepare the PEG/dextran blends, we used dextran from *Leuconostoc mesenteroides* with a molar mass of 450-650 kg/mol (Dextran500k; Sigma-Aldrich Japan, Tokyo, Japan), PEG with a molar mass of 6 kg/mol (PEG6k; FUJIFILM Wako Pure Chemical Co., Japan; Tokyo, Japan), and ultrapure distilled water (Invitrogen, CA; Catalog no. 10977-023). As a fluorescent labeled polymer, we used fluorescein-isothiocyanate-labeled dextran with molar masses of 500, 150, and 10 kg/mol (FITC-Dex500k, FITC-Dex150k, FITC-Dex10k; Sigma–Aldrich Japan), rhodamine-B-labeled dextran with a molar mass of 10 kg/mol (RB-Dex10k; Sigma–Aldrich Japan), and rhodamine-B-labeled PEG with a molar mass of 5 kg/mol (RB-PEG5k; Nanocs, MA, USA). Additionally, 1-Palmitoyl-2-oleoylsn-glycero-3-phosphocholine (PC) and a PEGylated lipid, 1,2-dioleoyl-sn-glycero-3-phosphoethanolamine-N-[methoxy(polyethylene glycol)-750] (ammonium salt) (PEG750-PE), were purchased from NOF Corporation (Tokyo, Japan) and Avanti Polar Lipids Inc. (AL, USA), respectively. Mineral oil (kinetic viscosity of 14–17 mm^2^ s^-1^ at 38 °C) was obtained from Nacalai Tesque (Kyoto, Japan) and treated using molecular sieves. Fluorescent molecule 5-carboxytetramethylrhodamine (TAMRA, Sigma–Aldrich Japan) was used for molecular diffusion analysis. These materials were used without further purification, except for mineral oil.

### Preparation of droplets encapsulated with lipid layer

Stock solutions of PEG6k and dextran500k with a concentration of 15wt %–20wt% were prepared by dissolving them in distilled water and then heating them to 60 °C. After allowing them to stand for 1 day at approximately 20 °C, the stock solutions were mixed to achieve the desired concentrations of PEG and dextran. For fluorescence observation, 0.1% FITC-Dex and 0.1% RB-PEG was added to the PEG and dextran solutions, respectively. The radius of gyration for PEG6k was ~3 nm, and the critical overlap concentration of dextran500k was ~4.7 wt%.^35^

Lipids in a mineral oil solution of ~1 mM PC (lipid-oil solution) were prepared by mixing a lipid-in-chloroform solution and mineral oil, followed by overnight evaporation at 60-70 °C^11^. We substituted 25 mol% PC into PEG750-PE for the PEGylated membrane system. W/O droplets encapsulated with a lipid layer were prepared as reported previously^18^. Briefly, the PEG/dextran blend in the 1-phase region was added to the lipid–oil solution at a volume ratio of 1:20 (W/O). By performing emulsification via pipetting, we prepared W/O droplets with radii *R* ranging from 5 to 70 μm. The droplet solution was injected into a custom-developed cover-slip cell comprising a ~0.1-mm-thick spacer (achieved using double-sided sticky tape).

### Fluorescence observation and intensity analysis

Fluorescence images of the bulk solutions and droplets were obtained using a confocal laser scanning microscope (IX83, FV1200; Olympus Inc., Tokyo, Japan) equipped with a water immersion objective lens (UPLSAPO 60XW, Olympus Inc.), or an inverted fluorescence microscope (IX71, 73; Olympus Inc., Tokyo, Japan) equipped with an oil immersion objective lens (UPLXAPO60XO, Olympus Inc.). FITC-Dex and RB-PEG were excited at wavelengths of 473 and 559 nm and detected in the ranges of 490–540 and 575–675 nm, respectively. The fluorescence intensity was analyzed using Fiji software (National Institute of Health (NIH), USA). In terms of the phase-separated droplets, we measured the average intensity of RB-PEG and FITC-Dex for the dextran-rich and PEG-rich phases, which yielded the *I*_max_ and *I*_min_ of RB-PEG (or FITC-Dex) for the PEG-rich and dextran-rich phases (or dextran-rich and PEG-rich phases), respectively. The intensity difference normalized by the background intensity *I*_0_, i.e., (*I*_max_ – *I*_min_)/*I*_0_, of RB-PEG and FITC-Dex was analyzed for the PEG-rich and dextran-rich phases, respectively. Subsequently, this value was used to evaluate the fractionation degree. To eliminate artifacts due to evaporation and fluorescence fading, fluorescence intensity measurements were performed within 0–120 min of droplet preparation.

### Determination of phase diagram of PEG/dextran blend

The 2-phase region of the PEG6k and dextran500k blends was identified primarily via turbidity confirmation and microscopic observation. To construct the binodal and tie lines, we prepared four different compositions with the same weight fraction of PEG and dextran (4 wt%, 4.5 wt%, 6 wt%, and 6.75 wt%). After centrifugation (5000 rpm for 60 min at ~20 °C; Model 3780; Kubota Co., Tokyo, Japan), the sample was allowed to stand for several days to reach thermal equilibrium. Subsequently, the polymer concentrations for the upper PEG-rich phase and lower dextran-rich phase were estimated by analyzing the refractive index (RI), density, and fluorescence intensity ratio. For (i, ii) RI and density analyses, we used a reflectometer (Abbemat 3000, Anton Paar GmbH, Graz, Austria) with an accuracy of 0.0001 at 589 nm and a density meter (DMA 1001, Anton Paar GmbH) with a resolution of 0.05 mg/mL. The values of density *ρ* and the RI indices *n* for the PEG and dextran solutions at 20 °C were plotted against the polymer concentration *w_i_*, as shown in Figures S1 (a, b). It was discovered that *ρ* exhibited a linear relationship with *w*, i.e., *ρ* = *ρ*_0_ + *Aw*. The fitting parameter of the solvent density *ρ*_0_ was 0.998, which is consistent with the literature value of *ρ*_0_ = 0.998 at 20 °C ^36^. The fitting parameter of *A* for PEG at 20 °C was *A* = 0.1728 mg/mL, which is slightly lower than that of *A* at 25 °C (i.e., *A* = 0.1752 mg/mL^37^). The value for dextran at 20 °C was *A* = 0.3767 mg/mL, which is similar to the previously reported value at 24 °C.^23^ The RI indexes *n* for PEG6k and dextran500k were plotted against *wρ*, as shown in Figures S1 (c, d), and fitted with *n* = *n*_0_ + (*dn/dc*)*wρ*, where *n*_0_ is the *n* for distilled water at 20 °C, i.e., 1.3 3 3.^36^ The values of (*dn/dc*) for PEG and dextran were 0.137 and 0.142 mL/g, respectively, which agree well with previously reported values^23^. We constructed binodal and tie lines based on both the relationship between *n* and (*dn*/*dc*), as well as the fluorescence intensity ratio normalized by the background, (*I*_max_ – *I*_min_)/*I*_0_, as mentioned above.

### Interfacial tension on surface of PC droplets

The interfacial tension on the surface of droplets encapsulated with a PC membrane was analyzed at approximately 20 °C using the pendant drop method (DM-501, Kyowa Interface Science Co., Saitama, Japan). Lipids in a mineral oil solution of ~1 mM PC were used as the oil phase. Because of the slow adsorption of lipid PC^29^ and the high viscosity of the dextran solution^15^, we analyzed the relaxation process for more than 15 min to obtain the saturation value.

### Diffusion analysis for 1-phase solution inside droplets

We analyzed the diffusion coefficient *D* of RB-PEG5k inside the PC droplets containing 5 wt% dextran and 1wt% PEG with 4 nM RB-PEG5k using point FCS analysis. FCS analysis was performed inside the droplet using a confocal microscope integrated with a molecular diffusion package (IX83, FV1200, Olympus Inc.). To minimize the RI mismatch and temperature change during laser irradiation, a water immersion objective lens (UPLSAPO 60XW) was used to perform point FCS analysis with minimal laser power for droplets that adhered to the cover glass, as reported previously.^15, 16, 17^ The illumination volumes were calibrated with TAMRA solution based on a diffusion coefficient of 280 μm^2^/s (Manual for diffusion analysis package, Olympus Inc.). RB-PEG5k and TAMRA were excited at a wavelength of 559 nm and detected within the 575-675 nm range. The software accompanying the microscope was used to calculate the autocorrelation and diffusion coefficients. The radius of gyration of RB-PEG5k was ~3 nm.

## Acknowledgements

We thank Mr. Kazuya Haganuma (University of Tokyo) for experimental supports. This research was funded by Japan Society for the Promotion of Science (JSPS) KAKENHI, grant number: 18H01187, 21H05871, 21K18596 (M.Y.), 20K14425 (C.W.), and Japan Science and Technology Agency (JST) ACT-X, grant number: JPMJAX191L (C. W.).

## Data availability

The authors declare that all the data supporting the findings of this study are available within the paper and its Supplementary Information.

## Contributions

M. Y. conceived the project. C. W., Y. K., T. F. and M. Y. did experiments to achieve the phase diagram in bulk in Fig. 1a. C. W., T. F., Y. K. and M. Y. performed experiments using droplets analyze the data detailed in Figs. 1cd, 2ab and 3ab. Y. K. and J. T. performed FCS experiments and Y. K., J. T. and M. Y. analyzed the data in Fig. 2c. C. W. conceived the experiments using PEGylated lipids detailed in Fig. 3cd. M. Y. conceived the experiments using shorter dextran detailed in Fig. 4. F. K., T. F., and M. Y. conceived the surface tension measurements detailed in Fig. 5a. M.Y. and Y. K. derived analyze the gain of droplet surface energy and depletion energy shown in Fig. 5b and Supplemental Information, S1. M. Y. and C. W. wrote the paper and the co-authors edited it.

