## Supplemental Information and Figures for "Competitive membrane wetting of polymer blends in artificial cells initiates phase separation and promotes fractionation"

AUTHOR INFORMATION

**Contents**

**S1. The estimation of depletion effect**

**Supplemental figures**

Figure S1. Refractive index and density of PEG6k and dextran500k solutions

Figure S2. Phase separation dynamics for 2 wt% PEG6k and 7 wt% dextran500k droplets

Figure S3. R-dependent LLPS inside Span 80 droplets containing 3 wt% PEG6k and 3 wt% dextran500k

### S1. The estimation of depletion effect

Under our experimental conditions using the PEG6k/dextran500k in 1-phase in bulk, the number of PEG6k is much larger than that of dextran500k. For example, the molar concentrations of 3 wt% PEG6k and 3 wt% dextran500k are approximately 5 mM and 60 nM, respectively. Assuming that PEG6k and dextran500k as small and large spherical colloids, the depletion energy gain when the two large colloids of dextran500k come into contact is expressed as follow based on Asakura-Oosawa theory[1]

$$\Delta F_{\text{gain}} \sim \left[1 + \left(\frac{3}{2}\frac{R_{\text{Dex}}}{R_{\text{Peg}}}\right)\right] n_{\text{Peg}} k_{\text{B}} T \quad \text{(S1)}$$

where $R_{\mathrm{i}}$ (i = Peg, Dex) are radii of PEG and dextran, $n_{\mathrm{Peg}}$ is the volume occupied by the PEG chains, $k_{\mathrm{B}}$ is the Boltzmann constant, and $T$ is the absolute temperature. As the $R_{\mathrm{i}}$ value, we use the radii of gyration for PEG and dextran with the molecular weight Mw as follows[2, 3],

$$R_{\text{Peg}} = 0.02 \times M_{\text{w}}^{0.58} \quad \text{(S2)}$$

$$R_{\text{Dex}} = 0.0633 \times M_{\text{w}}^{0.427}. \quad \text{(S3)}$$

The $R_{\mathrm{i}}$ for PEG6k, dextran500k and dextran10k is $R_{\mathrm{Peg6k}}$ ~3 nm, $\mathrm{R}_{\mathrm{Dex500k}}$ ~17.2 nm, and $R_{\mathrm{Dex10k}}$ ~3 nm respectively.

We estimate the depletion energy gain $\Delta F_{\mathrm{gain}}$ for 3 wt% PEG6k (far from the critical concentration of PEG6k, 8 wt%[2]) with a volume occupied by a PEG chain being $v_{\mathrm{Peg6k}}$. For simplicity, here we assume the maximum $n_{\mathrm{PEG}}$ based on the volume occupied by the PEG as $v_{\mathrm{Peg6k}}$ $\sim 4\pi R^3_{\mathrm{Peg6k}}/3$. For 3 wt% PEG6k, maximum $n_{\mathrm{PEG}}$ is approximately ~0.38. Substituting the values of $R_{\mathrm{PEG6k}}$, $R_{\mathrm{Dex500k}}$, and $n_{\mathrm{PEG}}$ into Eq. (S1), the depletion energy gain is derived as $\Delta F_{\mathrm{gain}}$ ~3.5 $k_{\mathrm{B}}$T. If the $n_{\mathrm{PEG}}$ is decreased to 0.3, the $\Delta F_{\mathrm{gain}}$ is ~2.8 $k_{\mathrm{B}}$T. Therefore, the free-energy gain via depletion interaction $\Delta F_{\mathrm{gain}}$ can be several $k_{\mathrm{B}}$T, which is larger than adhesion energy of a PEG chain to the lipid bilayer (~1.7 $k_{\mathrm{B}}$T),[4] a single H-bond (~1.5 $k_{\mathrm{B}}$T), and the van der Waals interaction (~0.1 $k_{\mathrm{B}}$T).[5]

The depletion layer exists near the droplet surface by the thickness of $R_{\mathrm{peg}}$ ~3.1 nm, the estimated depletion volume is sufficiently small compared to the volume of phase-separated droplets ($5 < R < 20$ μm). Therefore, such droplet size dependence on the depletion force will be negligible. Since the dextran500k has a wide range of Mw, the contained small dextran having Mw << 500k (dextran10k has a similar size with PEG6k) may contribute to the enhancement of the depletion interaction. If 1wt% component of 3 wt% dextran 500k is dextran10k, the molar mass is approximately 30 nM, which is much smaller than that of PEG6k, 5 mM. When estimating the effective depletion force from the local concentration of short PEG and dextran accumulated on the membrane, the depletion volume and small dextran contribution on the membrane must be considered.

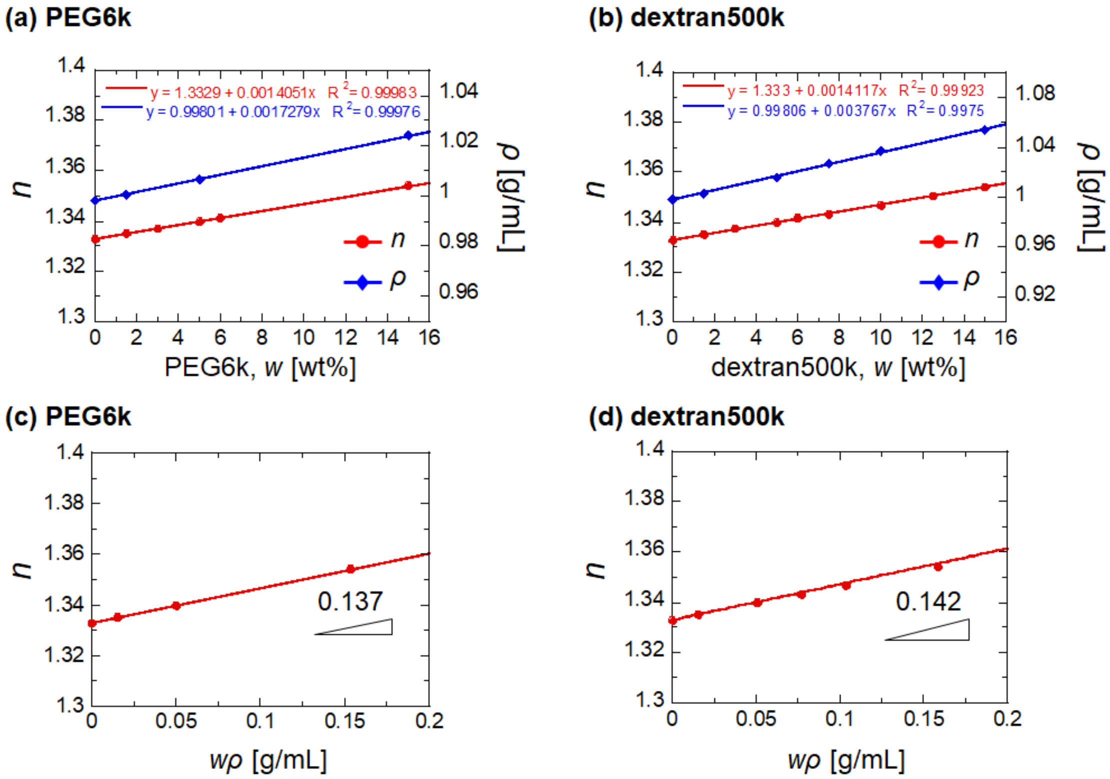


**Figure S1.** (a, b) Refractive index $n$ and density $\rho$ against various concentrations $w$ of PEG6k (a) and dextran500k (b). (c, d) Refractive index $n$ against $w\rho$ of PEG6k (c) and dextran500k (d).

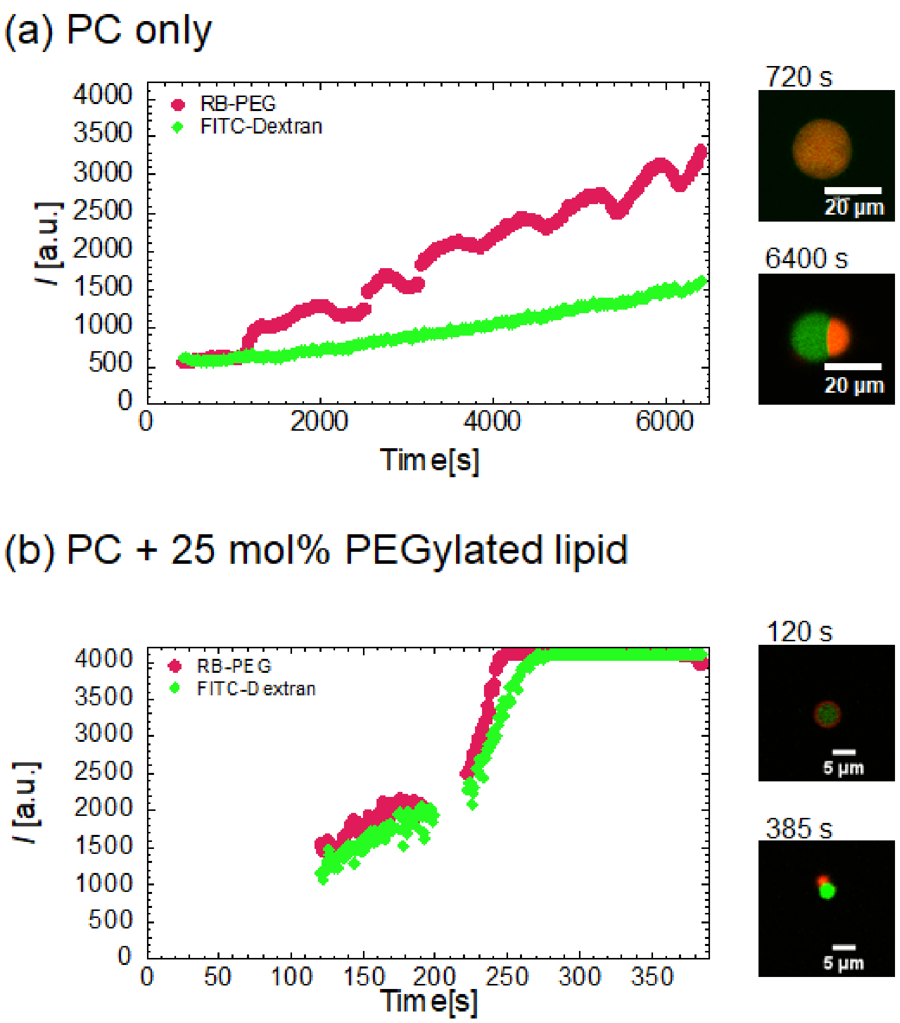


**Figure S2.** Phase separation dynamics for 2 wt% PEG6k and 7 wt% dextran500k droplets covered with a layer of (a) PC only (a) and (b) PC with 25 mol% PEGylated lipids. (left) Intensities of RB-PEG5k (green) and FITC-dextran500k (red) in PEG-rich phase and dextran-rich phase are plotted against time. (right) Fluorescence images of the droplets at the initial (upper) and late stages (lower) of phase separation, respectively.

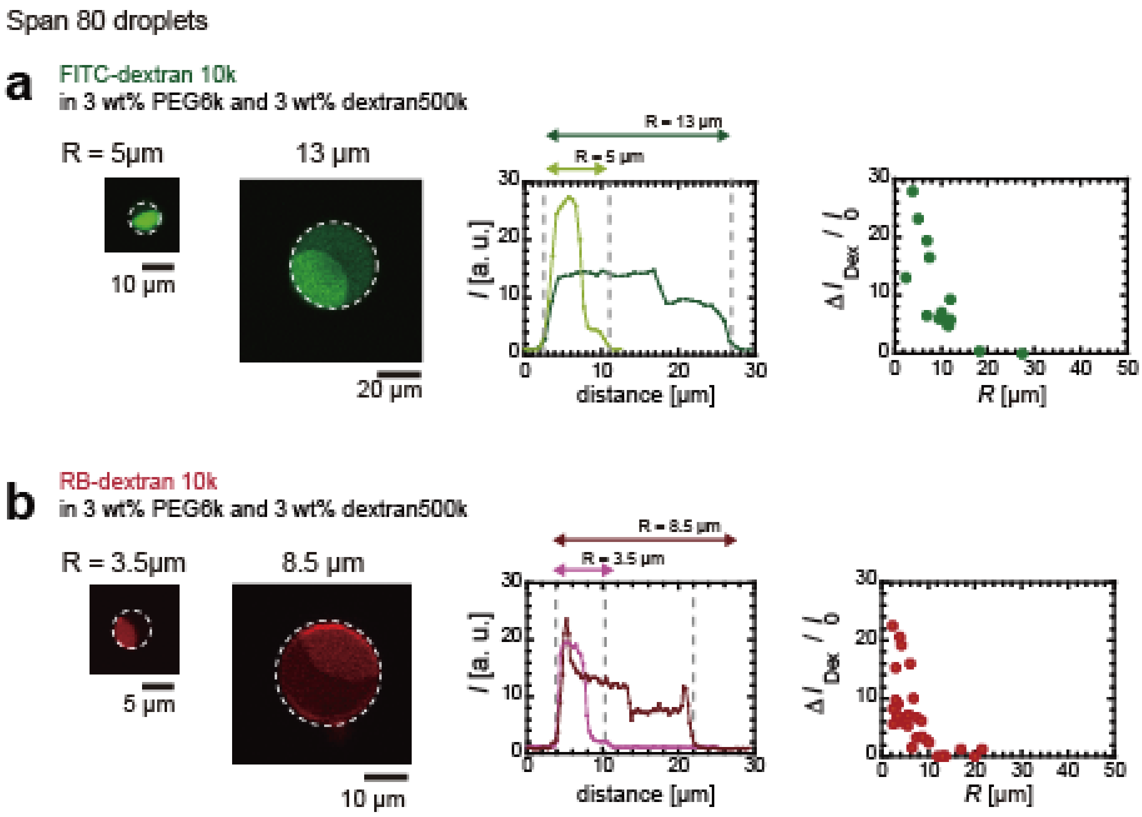


**Figure S3.** $R$-dependent LLPS inside Span 80 droplets containing 3 wt% PEG6k and 3 wt% dextran500k with (a) FITC-dextran10k or (b) RB-dextran10k. (left) Examples of confocal fluorescence images for phase-separated droplets with a different radius $R$ and their fluorescence intensity profiles along equational plane. The dashed lines indicate the position of the droplet surface. (right) Normalized intensity difference of FITC-dextran10k and RB-dextran10k, $\Delta I_{Dex}$ / $I_0$ plotted against $R$.
